# HSP90-CDC37-PP5 forms a structural platform for kinase dephosphorylation

**DOI:** 10.1101/2022.05.03.490524

**Authors:** Jasmeen Oberoi, Xavi Aran-Guiu, Emily A. Outwin, Pascale Schellenberger, Theodoros I. Roumeliotis, Jyoti S. Choudhary, Laurence H. Pearl

## Abstract

Activation of client protein kinases by the HSP90 molecular chaperone system is affected by phosphorylation at multiple sites on HSP90, on the kinase specific co-chaperone CDC37, and the kinase client itself. Removal of regulatory phosphorylation from client kinases and their release from the HSP90-CDC37 system depends on a Ser/Thr phosphatase PP5, which associates with HSP90 via its N-terminal TPR domain. Here we present the cryoEM structure of the oncogenic protein kinase client BRAF^V600E^ bound to HSP90-CDC37, showing how the V600E mutation favours BRAF association with HSP90-CDC37. Structures of HSP90-CDC37-BRAF^V600E^ complexes with PP5, in autoinhibited and activated conformations, together with proteomic analysis of its phosphatase activity, reveal how PP5 is activated by recruitment to HSP90 complexes to comprehensively dephosphorylate client proteins.

## INTRODUCTION

Interaction with the HSP90 molecular chaperone system is a prerequisite for the stability and biological function of a large proportion of the kinome ^1^, including most of the main oncogenic protein kinases ^2^. Recruitment of kinases to the HSP90 system is mediated by CDC37 ^3^, which functions as an adaptor able to interact independently with HSP90 and protein kinases, and facilitate their association ^4^. CDC37 is subject to a number of phosphorylation events ^5^, one of which – phosphorylation of Ser13 by casein kinase 2 (CK2) ^6,7^ – is critical to its function in protein kinase activation. HSP90 itself is also multiply phosphorylated ^8^, and while none are critical to its core biochemistry, several of the modified sites have nonetheless been shown to have important functions in regulation of ATP-utilisation and/or co-chaperone and client interactions ^9-13^. The protein kinase clients of HSP90-CDC37 are themselves frequently phosphorylated, sometime autogenously, as part of their regulation, and can in turn participate in phosphorylation of components of the chaperone complexes to which they are recruited, generating a complex network structure of post-translational regulation – the so-called Chaperone Code - the surface of which has only been scratched ^14^.

Phosphorylation is by its nature a reversible post-translational modification, and its role in switching the behaviour of a modified protein depends both on the kinase that ‘writes’ the modification and the phosphatase that ‘erases’ it ^15^. HSP90 is directly associated with an unusual serine/threonine protein phosphatase PP5 (Ppt1p in yeast), which has been implicated in the maturation/activation of a number of HSP90-dependent client proteins ^16-19^. PP5 has a tetratricopeptide (TPR) domain attached to the N-terminus of a Mn^2+^- dependent PP1/PP2A/PP2B family phosphatase domain ^20^. In common with several other HSP90-associated proteins, the TPR domain confers high affinity for the MEEVD motif that forms the extreme C-terminus of HSP90 ^21^. Ppt1p, the yeast homologue of PP5, has been implicated in regulating the phosphorylation of HSP90 itself, with deficit of Ppt1p activity leading to reduced activation of a range of client proteins *in vivo* ^22^. Within the specific context of the HSP90-CDC37 system, activation of protein kinase clients *in vivo* has been shown to depend on dephosphorylation of pSer-13 in CDC37 by PP5, which only occurs when PP5 and CDC37 are bound simultaneously to the same HSP90 dimer ^23^.

To understand how PP5 operates in the context of an HSP90-CDC37 client complex, we have reconstituted an active PP5 complex with HSP90, CDC37 and the highly HSP90-dependent V600E mutant of the protein kinase BRAF. We have determined the cryoEM structure of an HSP90-CDC37-BRAF^V600E^ complex, and structures of HSP90-CDC37-BRAF^V600E^ with PP5 bound in activated and autoinhibited conformations. These structures reveal how a single PP5 docks with the dimeric C-terminus of HSP90, and how docked PP5 rearranges to allow the catalytic phosphatase domain to access phosphorylation sites on the chaperone, co-chaperone and client. Together with proteomic analysis of PP5 activity on HSP90-CDC37-bound BRAF^V600E^ our studies reveal how the HSP90-CDC37-PP5 complex acts to comprehensively dephosphorylate the bound client.

## RESULTS

### PP5 dephosphorylates CDC37 within an HSP90-CDC37-BRAF^V600E^ complex

We previously showed that protein phosphatase 5 (PP5) was able to dephosphorylate pSer13 in CDC37 when both proteins were physically associated with HSP90 ^23^. To determine whether PP5 could also do this when CDC37 was engaged in a complex with a client protein and HSP90, we co-expressed and purified an HSP90-CDC37-BRAF^V600E^ (HCK) complex from insect cells (**see METHODS**), and used a phosphospecific antibody to demonstrate that wild-type PP5 could dephosphorylate CDC37-pSer13 in the complex in a time dependent manner (**SUPPL. FIG. 1A**).

To attempt to trap a productive complex of PP5 engaged with HSP90-CDC37-BRAF^V600E^, we incubated the HSP90-CDC37-BRAF^V600E^ complex with a PP5 D274N mutant which had previously been shown to catalytically inactivate PP5 with minimal disruption to substrate binding ^24^, and were able to purify a stable HSP90-CDC37-BRAF^V600E^-PP5 complex (HCK-P) on size exclusion chromatography (**SUPPL. FIG. 1B**).

### CryoEM structure determination

For structural studies the purified complex was cross-linked (**SUPPL. FIG. 1C**) and applied to cryoEM grids which were then plunged into liquid ethane. Movies from selected regions of the grids were recorded on an FEI Titan Krios microscope equipped with a Falcon IV detector (**see METHODS**). Movies were motion corrected, images processed, and particles picked using cryoSPARC ^25^and RELION 4.0 ^26^ (**SUPPL. FIG. 1D**,**E**). We obtained particle sets representative of three different structures which were separately refined. Final maps for HCK, HCK-P_open_, and HCK-P_closed_ complexes had overall resolutions of 3.4Å, 4.2Å, and 3.9Å respectively and allowed the fitting of substantive atomic models using the known crystal and cryoEM structures of the components (see **METHODS**). The image processing workflow and analysis of resolution are shown in **SUPPL. FIG. 2**,**3**.

### Structure of HSP90-CDC37-BRAF^V600E^

The structure of the HSP90-CDC37-BRAF^V600E^ complex consists of two molecules of human HSP90β arranged in the ATP-bound closed conformation originally observed in a complex of yeast HSP90 and the co-chaperone P23/Sba1 ^27^ (**FIGURE 1A**). The polypeptide chain for both HSP90β molecules can be traced through more or less continuous ordered density from Glu10 to Glu692, with the exception of the low-complexity ‘linker segment’ from approximately 220-275 which connects the N-terminal and central domains. Consistent with the closed conformation, bound ATP (or ADP-molybdate) is present in the N-terminal domains of both HSP90 molecules (**FIGURE 1B**).

**FIGURE 1.**
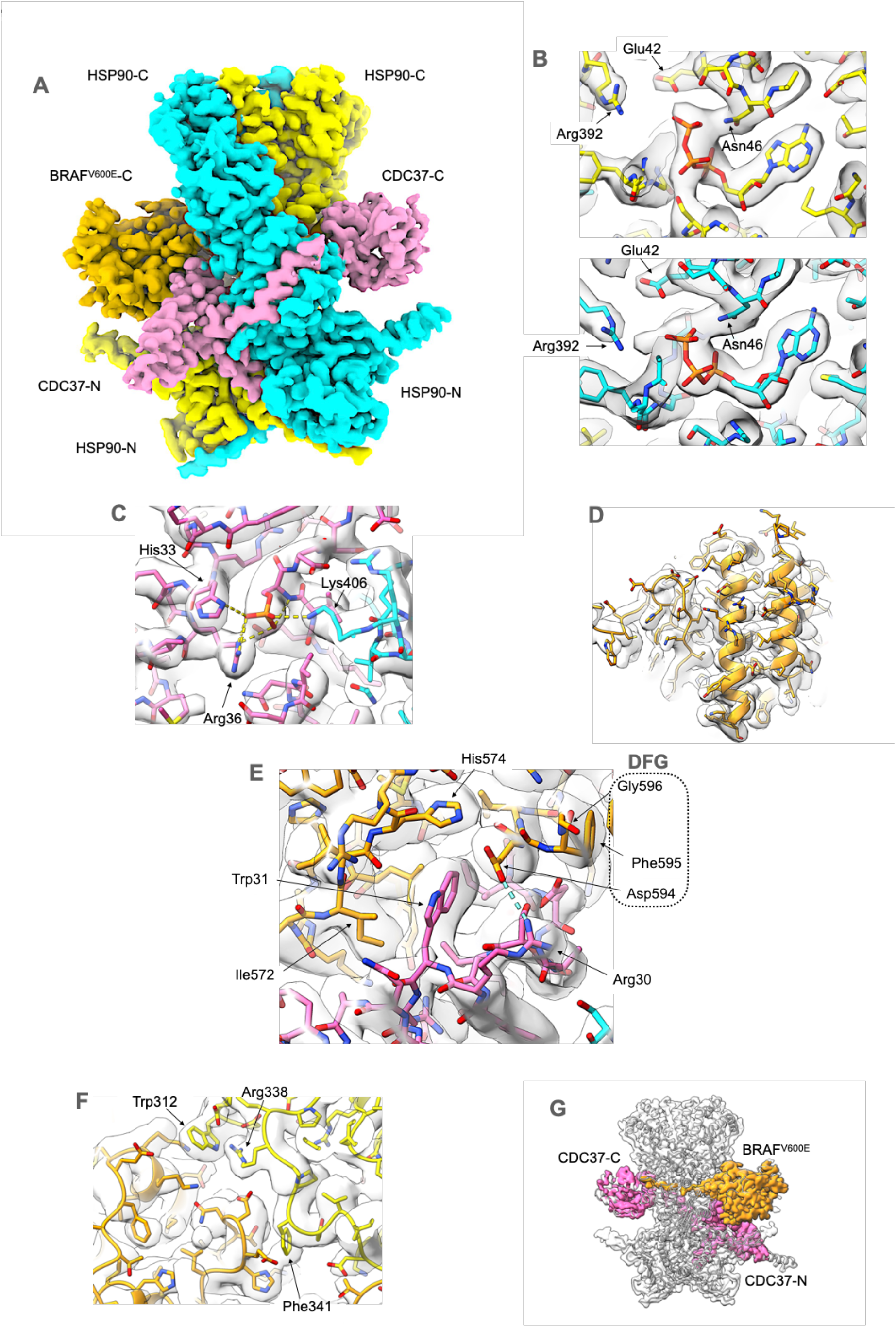
Structure of HSP90-CDC37-BRAF^V600E^ complex. **A**. Experimental cryoEM map of HSP90-CDC37-BRAF^V600E^ complex, surface coloured according to the underlying protein chain. The HSP90 dimer (blue and yellow – the colours of the Ukrainian flag) is in a closed conformation, with CDC37 (pink) wrapping around the edge of one of the HSP90 monomers. The C-lobe of the kinase domain of BRAF^V600E^ (orange) packs against the opposite face to the globular C-terminal half of CDC37, and interacts with the N-terminal coiled-coil α-hairpin of CDC37. This and all other molecular graphics were created using ChimeraX ^58^. **B**. ATP (or possibly ADP-molybdate) is bound in the N-terminal domain of both HSP90 monomers, and interacts with the catalytic Arg392 from the middle domain. **C**. Phosphorylated Ser13 of CDC37 interacts with His33 and Arg36 of CDC37, stabilising the conformation of the N-terminal part of CDC37 and bridging to Lys406 from the middle segment of one of the HSP90 monomers. **D**. The bound BRAF^V600E^ is well resolved throughout allowing for a nearly full tracing of its amino acid sequence in the cryoEM map. **E**. Within the complex, Trp31 displaces Phe595 of the key regulatory DFG motif into a different conformation than in the intact kinase, stabilised by interaction of DFG Asp594 with CDC37 Arg30. This conformational switch is facilitated by the oncogenic V600E mutation in the ‘activation segment’ immediately following the DFG motif, and explains why the oncogenic BRAF^V600E^ mutant is a strong client of the HSP90-CDC37 chaperone system whereas the wild-type is not ^32^. **F**. HSP90 itself only makes peripheral contact with the kinase C-lobe, but mutation of the HSP90 residues involved impair kinase activation *in vivo* ^35^. **G**. As previously seen for CDK4 in complex with HSP90 and CDC37 ^28^ the strand from the kinase N-lobe immediately upstream of the well-ordered C-lobe, becomes linearised and stretches between the two HSP90 monomers to emerge on the other face of the complex adjacent to the globular part of CDC37. No ordered structure upstream of this is visible in the cryoEM maps.

CDC37 in the complex presents in a very similar conformation as seen in the cryoEM structure of the HSP90-CDC37-CDK4 complex ^28^, with the N-terminus (residues 1-120) which consists predominantly of a long coiled-coil α-hairpin protruding from one side of the core HSP90 dimer, while the globular helical domain that forms the bulk of the C-terminal part (136-378) ^29^ is packed against the opposite face of the dimer (**FIGURE 1A**). The two halves of CDC37 are connected by an extended β-strand (121-135) which hydrogen bonds onto the edge of the central β-sheet of the middle domain of one of the HSP90 monomers. The polypeptide chain in the N-terminus can be traced from the N-terminal methionine to Cys54 and from Leu91 – Glu134, however the tip of the coiled-coil α-hairpin (residues 55-90) is not visible in the map. The C-terminal part of CDC37 is far less well defined than the N-terminus, with structure only discernible at the level of secondary structural elements from residue 140 to residue 266, suggesting a high degree of disorder and/or multiple conformational states for this loosely bound domain.

Serine 13, whose phosphorylation and targeted dephosphorylation are critical for client kinase activation by HSP90 ^6,7,23^, is clearly phosphorylated within the complex and engaged with the side chains of CDC37 residues His33 and Arg36, and Lys406 of HSP90 (**FIGURE 1C**).

Although the complex was formed by co-expression of the full-length proteins, relatively little of the 84kDa BRAF^V600E^ is visible in the cryoEM volume, with only the C-terminal lobe of the kinase domain being well defined in the map (**FIGURE 1D**). The polypeptide chain for this segment can be traced into clear features from Thr521 to Ile724, apart from the region corresponding to the ‘activation segment’ ^30^ connecting the 594-DFG-596 and 621-APE-623 motifs, which is poorly ordered. The final 42 residues at the C-terminus beyond the kinase C-lobe are also disordered.

The face of the BRAF^V600E^ kinase C-lobe that forms one wall of the ATP-binding cleft in the fully folded kinase structure ^31^ interacts with a contiguous segment of CDC37 from Thr19 to Ala35, incorporating the beginning of the first helix in the coiled-coil segment (**FIGURE 1E**). The core of the interface is provided by the side chains of His20, Ile23, Asp24, Ser27 and Trp31 of CDC37, which sit together in a channel in BRAF^V600E^ lined by Arg562, Gly563, Tyr566, Leu567, Ile572, His574, Thr590, Lys591, Ile592, Gly593 and Asp594.

One consequence of the interaction of CDC37-Trp31 with BRAF^V600E^ is to force the catalytically important DFG motif, into a quite different conformation to that found in the folded active kinase, with the following activation segment containing the oncogenic V600E mutated residue, being disordered. V600E and other common oncogenic Val600 mutations, have been shown to confer a strong dependence on association with HSP90-CDC37 for cellular stability and activation, whereas wild-type BRAF is a relatively weak ‘client’ ^32^.

Val600 in wild-type BRAF forms part of a hydrophobic cluster that holds the activation segment in an ordered inhibitory conformation ^33^, which is destabilised by oncogenic mutations such as V600E ^31,34^, contributing to unregulated kinase activity. Such destabilisation would also facilitate the conformational switch of the DFG and attached activation segment required by the interaction with CDC37 seen here, more readily than the hydrophobic and more rigid wild-type sequence, providing a satisfactory explanation for the substantially stronger HSP90-dependence of the oncogenic BRAF mutants.

HSP90 makes only a few direct contacts with the BRAF^V600E^ kinase C-lobe, restricted to peripheral interactions with surface exposed side chains of Arg338, Phe341 and Trp312 from the central region of HSP90 (**FIGURE 1F**) – the latter two previously implicated in client interactions in an earlier mutagenesis study ^35^, and a polar interaction between HSP90-Arg196 and BRAF-Asp565. The major interactions between HSP90 and the kinase client, involves residues 521-533 of BRAF^V600E^, which would be part of the N-terminal lobe in the fully folded kinase structure, which in the complex threads between the central segment of the two HSP90 monomers adjacent to the extended loops from Asn351 to Phe344 that come close together at the heart of the HSP90 dimer (**FIGURE 1G**). Upstream of BRAF^V600E^ residue 521, the chain emerges on the opposite face of the dimer, to run adjacent to the loosely bound globular domain in the C-terminal half of CDC37. However, the map in this region lacks detail due to conformational flexibility, and/or the presence of multiple conformations.

### Structure of HSP90-CDC37-BRAF^V600E^-PP5 Complexes

Two different sets of particles were obtained in which additional volume corresponding to PP5 was evident bound to the C-terminus of the HSP90 dimer within the HSP90-CDC37-BRAF^V600E^ complex. The two structures are distinguished by whether the C-terminal phosphatase domain of PP5 is packed against the N-terminal TPR domain in a ‘closed’ conformation or is substantially separated from it in an ‘open’ conformation. The locations and conformation of the HSP90 monomers and the visible parts of CDC37 and BRAF^V600E^ are essentially the same as in the HSP90-CDC37-BRAF^V600E^ complex (see above).

In the PP5-closed conformation (**FIGURE 2A**), the convex face of the TPR domain is juxtaposed with the active site of the phosphatase domain in a similar manner to that previously observed in the crystal structure of an autoinhibited conformation of PP5 ^36^ (**FIGURE 2B**). The concave face of the TPR domain is directed away from the phosphatase domain, and the channel formed by the TPR helices on this side of the domain is occupied by a feature consistent with the bound C-terminal -MEEVD peptide of HSP90, when compared to the NMR structure ^21^ (**FIGURE 2C**). However, the segment linking this to the globular core of HSP90 is not visible. An additional interface with HSP90 is made by the distal end of the last α-helix of the PP5-TPR domain centred on PP5-Phe148, which packs into a hydrophobic pocket at the C-terminus of the HSP90 dimer close to the dimer interface, lined by Ile684, Gly687 and Leu688 from the HSP90 monomer most likely providing the interacting MEEVD peptide, and Ala650, Asp653, and Leu654 of the other HSP90 monomer (**FIGURE 2D**).

**FIGURE 2.**
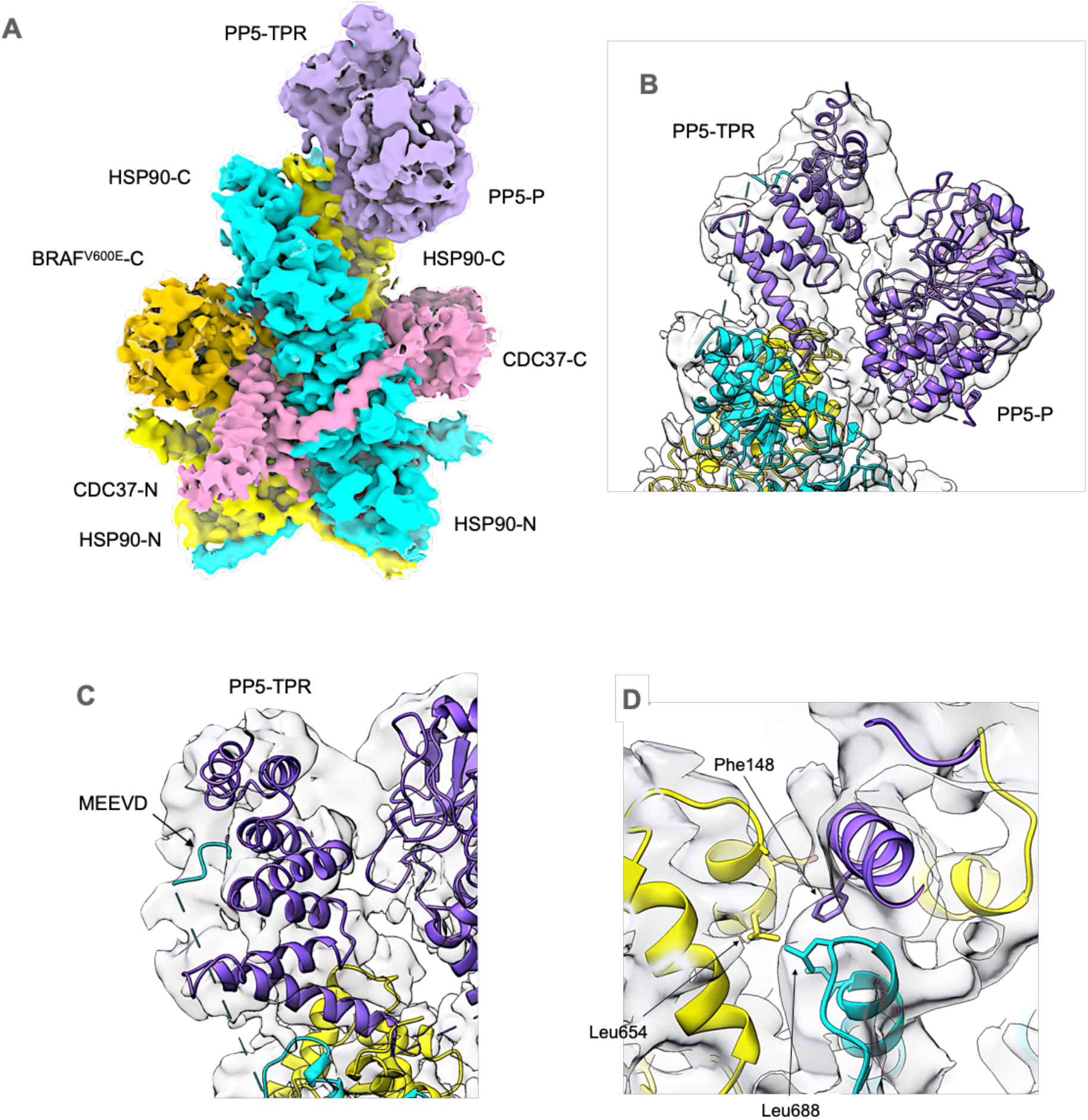
Structure of HSP90-CDC37-BRAF^V600E^ complex with autoinhibited PP5. **A**. Experimental cryoEM map of HSP90-CDC37-BRAF^V600E^-PP5 complex, surface coloured as **FIG. 1**, but with the addition of PP5 (purple), bound to the C-terminal end of the HSP90 dimer. **B**. PP5 is bound in the auto-inhibited closed conformation ^36^ in which the convex face of the TPR domain (PP5-TPR) occludes the substrate binding cleft of the phosphatase domain (PP5-P). As in previous PP5 crystal structures, the flexible linker connecting the last elongated α-helix of the TPR domain and the start of the phosphatase domain, is disordered. **C**. Although the resolution is insufficient for direct modelling, superimposition of NMR structures of the isolated PP5-TPR domain in complex with HSP90 C-terminal peptides ^21^ on the cryoEM volume, indicates the presence of the C-terminal MEEVD sequence bound in the convex face of the TPR domain. **D**. Additional to the HSP90-MEEVD interaction, the last helix of the PP5-TPR domain packs against a cluster of helices from the C-terminal domains of both HSP90 monomers, with the side chain of Phe148 at the tip of the last PP5 helix in a hydrophobic pocket.

In the PP5-open conformation (**FIGURE 3A**), the two domains of PP5 are completely separated, with the C-terminus of the TPR domain at Arg150 more than 50Å away from the N-terminus of the globular phosphatase domain at Tyr176. As with the closed complex, the cryoEM map suggests the presence of the HSP90 C-terminal MEEVD peptide bound into the concave face of the TPR, and the terminal helix of the TPR domain makes the equivalent interaction with the hydrophobic pocket formed by the C-terminal domains of the HSP90 dimer (**FIGURE 3B**).

**FIGURE 3.**
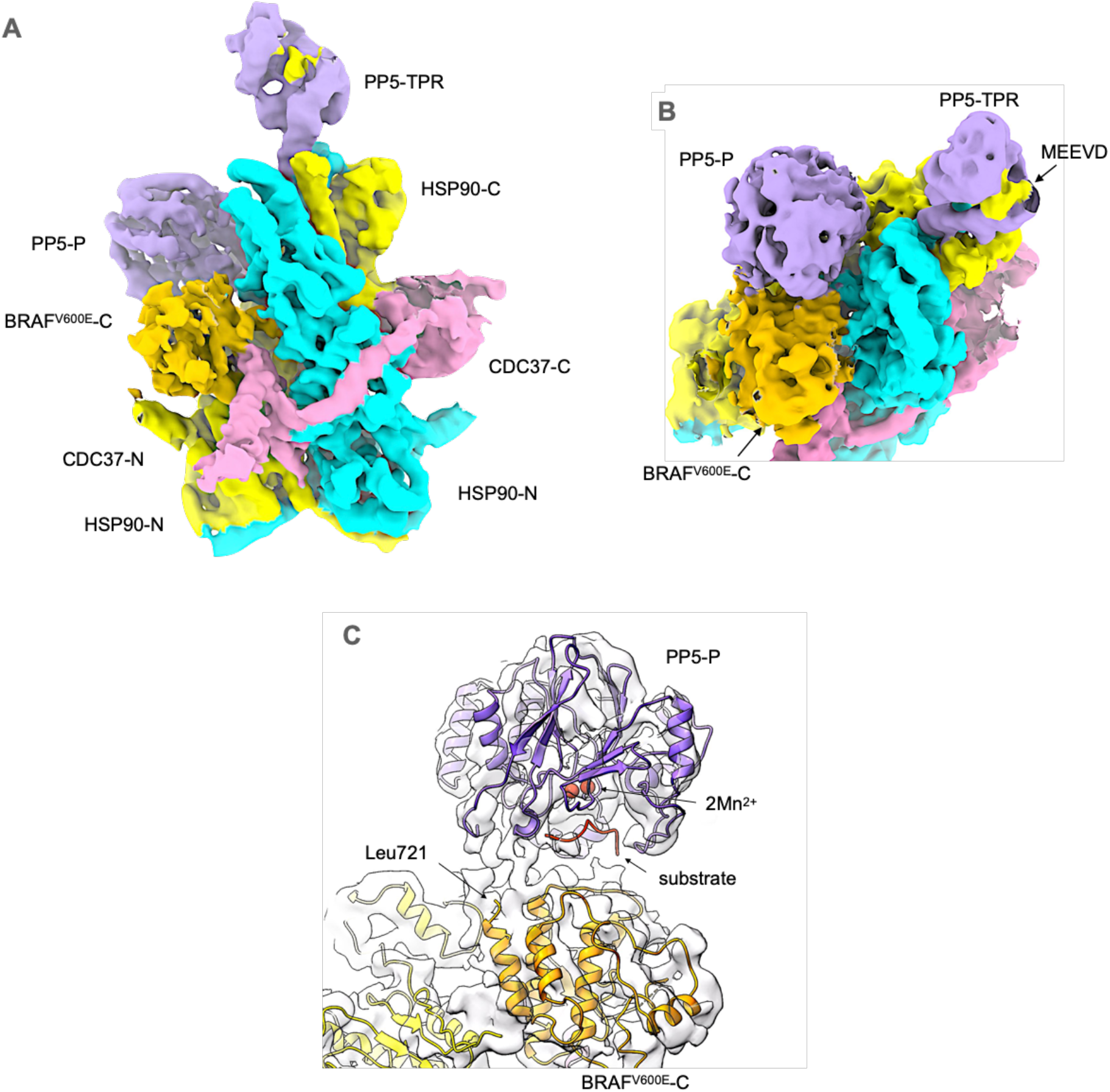
Structure of HSP90-CDC37-BRAF^V600E^ complex with activated PP5. **A**. Experimental cryoEM map of of HSP90-CDC37-BRAF^V600E^-PP5 complex, surface coloured as **FIG. 2**. **B**. Like the autoinhibited complex, the cryoEM map shows evidence of a bound C-terminal HSP90 MEEVD motif, and the tip of the last helix of the TPR packs into the equivalent hydrophobic pocket in the C-terminal domains of HSP90. However, the phosphatase domain of PP5 is no longer packed against its own TPR domain, but is instead engaged with the ordered kinase C-lobe of BRAF^V600E^. **C**. PP5 phosphatase domain is docked against the kinase C-lobe of BRAF^V600E^ with its substrate-binding cleft facing the inter-helix loops. The end of the last α-helix in the BRAF^V600E^ C-lobe at Leu721 is well positioned for phosphorylated residues downstream (including Ser729 – see **FIG 4**.) to be engaged productively with the phosphatase active site. The positions of a bound substrate peptide and the catalytic manganese ions of PP5 are modelled from 5HPE ^24^.

The detached phosphatase domain binds down towards the middle of the complex, bridging between surface loops at 461-467 in the middle domain of one HSP90 monomer and 569-574 in the C-terminal domain of the other monomer. In this position, the substrate-binding cleft of the phosphatase domain is in direct contact with the HSP90-bound C-lobe of the BRAF^V600E^ close to several inter-helix loops, and the point at which the unstructured C-terminus of BRAF^V600E^ would extend from the globular C-lobe. The early part of this BRAF^V600E^ segment downstream of residue 721 would be well positioned to interact productively with the phosphatase active site (**FIGURE 3C**).

As HSP90 is dimeric, there are two symmetrical disposed copies of the C-terminal hydrophobic binding site that the PP5 TPR domain interacts with. Binding of CDC37 and the kinase client render the overall complex asymmetrical, but as these are bound by the middle domain, the two-fold symmetry of the HSP90 C-terminus is largely unaffected. However, the two sites are sufficiently close together that binding of PP5 to one site sterically occludes the other, thereby restricting the stoichiometry to a single PP5 per HSP90 complex.

Fascinatingly, the PP5 TPR domain in the closed complex binds to one site such that the phosphatase domain is held on the face of the HSP90 dimer that presents the globular domain of CDC37, while in the open complex the TPR binds to the symmetry equivalent site so that the phosphatase domain is on the face that presents the kinase C-lobe and the coiled-coil helical hairpin of CDC37.

### PP5 Phosphatase Targets

While dephosphorylation of CDC37-pSer13 is the best studied HSP90-associated activity of PP5 ^23^ (and see **SUPPL. FIG 1A**), under the conditions in which HSP90-CDC37-BRAF^V600E^ is expressed and purified to be amenable to structural studies, CDC37-pSer13 is fully buried in the core of the ATP-bound closed HSP90 complex and remains so in the presence of the catalytically dead PP5. Even though the phosphatase domain of PP5 can detach from the C-terminus of the HSP90 dimer and move substantially towards CDC37, pSer13 would only become accessible when the N-terminal domains of HSP90 separate following ATP hydrolysis, so trapping a structurally tractable complex in which PP5 is engaged with CDC37-pSer13 remains to be achieved.

However, CDC37 is not the only component of the complex that is susceptible to phosphorylation, and therefore a potential substrate for PP5. To gain some insight into other potential substrates, we mapped the phosphorylation sites on the purified HSP90-CDC37-BRAF^V600E^ complex with and without treatment with PP5, by mass spectrometry (see **METHODS, SUPPL. FIG. 4, FIGURE 4A**). We identified two sites in HSP90 (Ser226, Ser255) which were significantly diminished by PP5 treatment. Both of these CK2 sites are within the charged linker segment connecting the N and middle domains of HSP90, and have been implicated in regulation of HSP90β secretion ^37^. We identified 12 sites in BRAF^V600E^ whose phosphorylation was significantly (p < 0.05) decreased by PP5 treatment (see **SUPPLEMENTARY MATERIAL**). One (Ser151) occurs just before the RAS binding domain (RBD), while six (Ser339, Ser365, Thr401, Ser429, Ser432, Ser446) occur within the disordered segment between the RBD and kinase domains. Ser365 plays a critical role in 14-3-3 binding ^33^ and along with Ser429 has been shown to have differential regulatory effects on different BRAF isoforms ^38^, while Ser446, which maps just upstream of the kinase N-lobe, is the topological equivalent of Ser338 in CRAF whose dephosphorylation by PP5 was previously shown to deactivate kinase signalling activity ^16^.

**FIGURE 4.**
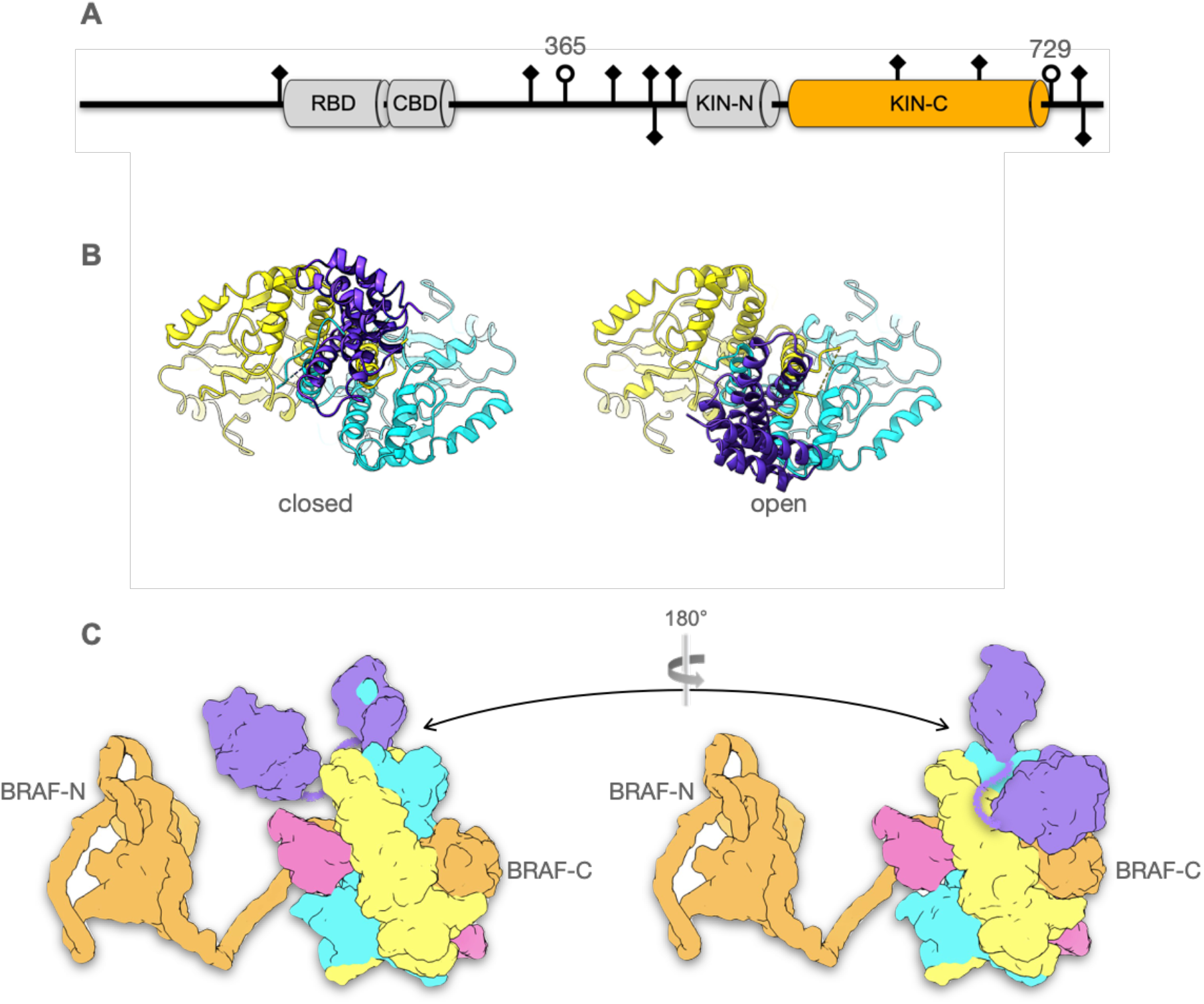
PP5 is a comprehensive remover of client phosphorylations. **A**. Schematic of phosphorylation sites identified in HSP90-CDC37-associated BRAF^V600E^ that are removed by addition of PP5 (see METHODS). Of the characterised globular regions, only the kinase C-lobe is ordered in complexes with HSP90-CDC37. Regulatory roles have been assigned in the literature for most of the sites identified. Of particular interest are 365 and 729 (open circles) as they mediate the interaction of BRAF in an autoinhibited complex with 14-3-3 proteins ^33^. Based on the relative position of the C-lobe and PP5-phosphatase, pSer729 is most likely to be the residue bound at the active site of the catalytically inactivated PP5 in the complex structure. **B**. The HSP90 dimer provides two symmetry equivalent alternative and mutually exclusive binding sites for the TPR domain of PP5, which are both used in the complexes, directing the associated phosphatase domain to different faces of the complex. **C**. By switching PP5 binding between the two TPR-binding sites on the HSP90 dimer, the flexibly attached phosphatase domain can access phosphorylation sites upstream and downstream of the core interacting domain of the kinase client, taking advantage of the partly unfolded and ‘linearised’ state that binding to HSP90-CDC37 facilitates.

Within the kinase domain itself, which is the focus of interaction of the HSP90-CDC37 system, we identified no phosphorylation sites in the N-lobe, but two sites within the C-lobe, which is the only part of BRAF^V600E^ in complex with HSP90-CDC37 that is resolved in the cryoEM structure. Ser614, identified as an inhibitory phosphorylation specifically enriched in the V600E mutant ^39^, maps to the C-terminal end of the activation segment 594-623 which is disordered in the structure, while Ser675 involved in regulation of BRAF ubiquitylation by the E3 ligase ITCH ^40^ is in the middle of an extended coil that connects two helices. Both of these residues map to parts of the surface of BRAF^V600E^ that are not involved in interaction with HSP90 or CDC37, and would therefore be accessible to the PP5 phosphatase domain given the flexibility of its connection to its TPR anchor. We found no phosphorylation of the activation segment residues Thr599 and Ser602 whose phosphorylation by MLK3 ^41^ is required for full BRAF (wt or V600E) kinase activity ^42^ suggesting that this occurs after release of the kinase from the chaperone complex.

Three further sites (Ser729, Ser750 and Thr753) all occur in the unstructured segment that follows the end of the C-lobe at residue 720. Ser729 has recently been shown to have a key role in 14-3-3 binding in concert with Ser365 ^33,43^. The proximity of the C-terminal end of the C-lobe to the substrate binding cleft of the phosphatase domain strongly suggests that one or more of these sites are engaged with the catalytically inactivated PP5.

## DISCUSSION

The structures presented here provide a view of a protein kinase other than CDK4 ^28^ in a complex with CDC37 and HSP90 in the ATP-bound closed state ^27^. This confirms a common mechanism of partial denaturation of the kinase domain, with the first strand of the N-lobe linearised in the molecular clamp of the closed HSP90. While the remainder of the N-lobe of CDK4 was partially visible in some subsets of particles, the smaller and less structured N-lobe of BRAF is completely disordered in the HSP90-CDC37-BRAF^V600E^ complex. The molecular details of the interaction of the DFG motif of BRAF^V600E^ with CDC37 in the complex provide a satisfactory explanation for how the oncogenic mutation of Val600 within the early part of the activation loop, converts BRAF from a weakly dependent HSP90 client with only moderate affinity for CDC37 ^1^, into a highly dependent client which is rapidly degraded when cells are treated with HSP90 inhibitors ^32,44^.

PP5 docks onto HSP90-CDC37-BRAF^V600E^ through a bipartite interaction mediated by the TPR domain, which binds the C-terminal MEEVD motif of HSP90 in its concave channel and plugs the tip of its terminal α-helix into one of two hydrophobic pockets formed at the interface of non-equivalent α-helices from each of the two HSP90 monomers. This mode of interaction is markedly different from that of FKBP51 with HSP90, which uses an N-terminal extension to its TPR to bind perpendicularly between the last helices of the HSP90 dimer ^45^. We observe PP5 binding alternatively to both C-terminal pockets on HSP90, but with markedly different conformations depending on which side of the overall complex the phosphatase domain is positioned (**FIGURE 4B**). When on the same face as the globular region of CDC37, which presents no phosphorylated substrate residues, the phosphatase domain remains associated with the TPR domain at the C-terminus of the HSP90 dimer in an auto-inhibited conformation ^36^. However, when bound with the phosphatase on the same face as the ordered C-lobe of the kinase, the phosphatase detaches from the TPR and docks against the middle domain of HSP90, with its substrate binding cleft in contact with the face of the kinase C-lobe from which the C-terminal segment extends, most likely held there by its interaction with one of the substrate phosphorylation sites that map to the early part of that segment. Considerable flexibility of the unstructured linker that connects the phosphatase to its TPR anchor, would allow the phosphatase access to other substrate phosphorylation sites on the exposed surface of the C-lobe, and indeed to parts of the kinase that are disordered in the complex, but nonetheless brought into general proximity to the phosphatase domain by their mutual binding to the HSP90-CDC37 ‘scaffold’.

### HSP90-CDC37-PP5 – a workbench for cleaning kinases

HSP90 sits at the heart of signal transduction within the eukaryotic cell ^46^, a substantial proportion of which is mediated by reversible phosphorylation. HSP90 in concert with its kinase-specific targeting partner CDC37, plays a critical role in the activation of the protein kinases that mediate this signalling, but the precise nature of that role remains obscure.

Biochemical ^4,47^ and structural studies ^28^ (and see above) have clearly shown that interaction with the HSP90-CDC37 system results in catalytic silencing of a client kinase, through partial unfolding of the kinase domain. However, association with HSP90-CDC37 also brings client kinases into the proximity of HSP90 co-chaperones that interact with the C-terminal MEEVD motif of the chaperone via their TPR domains. This exposes the client to modifications delivered by the catalytic domains of the TPR-cochaperones, which range from prolyl isomerases ^45,48^, to E3-ubiquitin ligases ^49,50^, and of most significance for protein kinase clients, a protein phosphatase - PP5.

Our data show that the overwhelming majority of Ser/Thr phosphorylations present on BRAF^V600E^ bound to HSP90-CDC37, are removed by PP5. Thus, PP5 effectively provides a ‘factory reset’ of the client kinase by removing whatever regulatory modifications may have been applied to it before it bound to HSP90-CDC37, on both N- and C -terminal sides of the kinase domain that drives chaperone recruitment (**FIGURE 4C**). Together with its ability to also remove the phosphorylation of CDC37 and thereby destabilise the association of the kinase client with the chaperone complex ^23^, PP5 provides a directionality to the process, ensuring the release of the client from the HSP90-CDC37 platform as a *tabula rasa*, ready for whatever new phosphorylations are required for its regulated function in the cell.

## Supporting information

Mass Spectrometry Data

## ACKNOWLEDGMENTS

We thank Rebecca Thompson, Emma Hesketh and Louie Aspinall for assistance with cryoEM data collection at The University of Leeds, Fabienne Beuron for assistance and advice regarding grid preparation, Basil Greber for advice on image processing, Antony Oliver for assistance with data handling, Lihong Zhou for assistance with insect cell expression, and Cara Vaughan and Chris Prodromou for contributions at earlier stages of the work. EM facilities at the Astbury Biostructure Laboratory at the University of Leeds are funded by University of Leeds ABSL award and Wellcome Trust award 108466/Z/15/Z. EM facilities at Sussex University are funded by Wellcome Trust Award Enhancement Grant 095605/Z/11/A – to L.H.P. - and the RM Phillips Trust. This work was supported by a Wellcome Trust Investigator Award 210719/Z/18/Z (L.H.P., J.O., X.A.G., E.A.O.) and Cancer Research UK Centre Grant C309/A25144 (T.I.R and J.S.C).

## AUTHOR CONTRIBUTIONS

Conceptualization :J.O, L.H.P.; Methodology : J.O., X.A.G., E.A.O., P.S., T.I.R., J.S.C., L.H.P.; Validation : J.O., J.S.C., L.H.P.; Formal Analysis : J.O., T.I.R., J.S.C., L.H.P.; Investigation : All Authors; Writing – Original Draft : L.H.P.; Writing – Review & Editing : All Authors; Visualisation L.H.P.; Supervision : L.H.P.; Funding Acquisition : L.H.P.

## COMPETING INTERESTS

The authors declare no competing interests.

## METHODS

### Protein expression and purification

Full-length human HSP90β, CDC37 and BRAF^V600E^ were subcloned into the baculovirus vector pBIG1a ^51^ with an N-terminal His^8^ tag on Hsp90β, a C-terminal His^8^ on CDC37 and N-terminal His^8^-2xStrep tag on the Braf^V600E^. Human rhinovirus 3C protease recognition sites were introduced between the proteins and the fusion tags.

*Sf9* cells were transfected with 1 µg of pBIG1a HSP90β, CDC37 and BRAF^V600E^ for viral production. For protein expression, *Sf9* cells were infected with HSP90β, CDC37 and BRAF^V600E^ baculovirus at a MOI of 2 and incubated for 72 h at 26 °C.

*Sf9* cells were lysed and sonicated in 40mM Hepes pH 7.4, 150mM Nacl, 10mM KCl, 20mM Na_2_MoO_4,_ 20mM imidazole, 0.5mM TCEP, 10% glycerol, 2U/ml Turbo DNAase (Invitrogen), EDTA-free protease inhibitor cocktail tablets and phosphatase inhibitor tablets (Roche). The NaCl concentration was increased to 750mM before incubating the lysate with talon resin (Takara Bio) for 2 hours at 4_°_C. The resin was washed sequentially with lysis buffer containing 750-600-450-300 and 150 mM NaCl. The protein complex was eluted from the resin in 40mM Hepes pH 7.5, 150mM NaCl, 10mM KCl, 20mM Na_2_MoO_4_, 500mM imidazole, 0.5mM TCEP, 10% glycerol. The eluate from the Talon resin was applied to a 2ml streptactin column (IBA) in Streptactin binding buffer consisting of 40mM Hepes pH 7.4, 150mM NaCl, 10 mM KCl, 10mM MgCl2, 20mM Na_2_MoO_4_, 0.5mM TCEP, 10% glycerol and eluted in binding buffer with 75mM Biotin. Elutions from the Streptactin column were applied to a Superdex 200 26/60 size exclusion column (GE Healthcare) and eluted in Streptactin binding buffer.

Human PP5 residues 16-499 with or without a D274N mutation were cloned into pGEX6P1 with an N-terminal GST tag and C-terminal His_6_ tag. PP5 was expressed in E.coli and purified as previously described _36_.

### HSP90-CDC37-BRAF^V600E^-PP5 complex assembly

To assemble the PP5 complex for CryoEM, the HSP90-CDC37-BRAF^V600E^ complex was purified as described above, but after the complex was eluted from the Strep-tactin column the Na_2_MoO_4_ and biotin were removed from the buffer by buffer exchanging using a 100kDa Mwt cut-off concentrator. This sample was then incubated with a 2 x molar excess of PP5 for 2 hours at 4^°^C. The sample was loaded onto a Superdex s200 10/300 size exclusion column (GE Healthcare) and eluted in 100mM NaCl, 25mM Hepes, 10mM KCl, 1mM MnCl_2_, 0.2mM TCEP and 2% glycerol. Samples prepared for CryoEM were further crosslinked with 1mM BS3 (Fisher Scientific UK Ltd) for 30 minutes at room temperature and quenched with 20mM Tris pH 7.5.

### CryoEM grid preparation data collection

Prior to grid preparation, the crosslinked HSP90-CDC37-BRAF^V600E^-PP5 complex was concentrated to 1μM and 3μl of the sample was applied to carbon grids (Quantifoil R1.2/1.3, Cu, 300 mesh) which were glow discharged using a Tergeo Plasma Cleaner (Pie Scientific). The sample was blotted for 5 seconds using a Leica EM GP2 (Leica microsystems) and plunge frozen in an ethane:propane mixture.

Three different datasets were collected using a Titan Krios (Thermo Fisher Scientific) equipped with Falcon 4 camera in counting mode, at a magnification of 96,000, which corresponds to a pixel size of 0.86 Å/pixel. EPU software version 2.11 (Thermo Fisher Scientific) was used to collect data with a defocus range of −2.5 to −1.3 μm at a dose rate of 9.5 e^-^/Å^2^/s, for a total exposure of 4.7 seconds and with 56 frames resulting in a total dose of 45 e^-^/Å^2^.

### CryoEM data processing

Movies of images from the three datasets were motion corrected separately using MotionCor2 ^52^ and were binned to 1.72 Å/pixel. A total of 21,196 micrographs were collected from the three different datasets. CTF estimation, particle picking and a first round of reference free 2D classification was carried out on each dataset separately initially. CTF was estimated using Patch CTF in cryoSPARC2 v3.3.1 ^53^. About 500 particles were manually picked initially to generate 2D templates for picking using Topaz ^54^. One round of reference-free 2D classification of particles picked using Topaz was performed in CryoSPARC to remove noisy class averages. At this stage the particles from the first round of 2D classification from each dataset were combined and a second round of 2D classification was performed on the combined total of 721,941 particles. Class averages with high resolution features were selected after this round of 2D classification and three Ab initio models were generated from 537,780 particles using CryoSPARC.

All subsequent processing steps were done in RELION4 ^55^. An initial round of 3D classification was performed using the particles and the ab-initio model obtained in CryoSPARC. Three classes had clearly recognizable density for HSP90, CDC37, BRAF and PP5 and were subjected to a second round of 3D classification. After this round, three distinct classes were observed, one containing only HSP90, CDC37 and BRAF, one containing HSP90, CDC37, BRAF and PP5 which is a more closed conformation bound to the C-terminal of HSP90 and the third containing HSP90, CDC37, BRAF and PP5 which is opened up and the TPR domain is bound to the C-terminal of HSP90 but where the phosphatase is engaged in the middle domain region of HSP90. To use the best signal from the HSP90-CDC37-BRAF complex class for particle polishing, particles from the three classes were combined (533,127 particles in total) and refined to 3.6 Å resolution. Only HSP90-CDC37-BRAF density was visible in this class. Particle polishing and CTF refinement was performed on these particles, followed by 3D refinement. A round of 3D classification was performed to retrieve back the three classes which now contain polished particles. The overall resolution for all three classes improved after 3D refinement.

To improve the resolution of the PP5 domains, the particles from the two classes which contained PP5 were further classified using signal subtraction and focused 3D classification without alignments, with masks around the PP5 domains. After reverting to original particles and applying a soft mask around the whole complex, the best focused class for the HSP90, CDC37, BRAF and PP5 (in a more ‘closed’ conformation) refined to a resolution of 3.9 Å and the best class for the HSP90, CDC37, BRAF and PP5 (in an open conformation) refined to 4.2 Å, with improved density observed for the PP5 domains in both classes. Particles from all three classes were re-extracted at 0.86A/pix and a final round of particle polishing was performed on all three classes individually, followed by postprocessing in RELION in which the nominal resolution was determined by the gold standard Fourier shell correlation (FSC) method ^56^. Maps were subsequently post-processed using DeepEMhancer ^57^.

### Model Building

Atomic models were derived from the cryoEM structure of an HSP90-CDC37-CDK4 complex (PDB code: 5FWK), and crystal structures of PP5 protein and domains (PDB codes : 1WAO, 5HPE and 1A17) and BRAF kinase domain (PDB code: 1UWH) docked as rigid bodies into experimental volumes using ChimeraX ^58^. The local fit of the models was adjusted manually in Coot ^59^ and the global fit optimised using phenix.refine ^60^. Parameters defining the data collection and the quality of the final atomic models are given in **SUPPLEMENTARY TABLE 1**. Models and maps have been deposited in PDB and EMD as follows : HCK - PDB ID 7ZR0, EMD-14875; HCKPo - PDB ID 7ZR6, EMD-14884; HCKPc PDB ID 7ZR5, EMD-14883.

### Dephosphorylation Assays

To monitor dephosphorylation of CDC37-pSer13 by PP5, 0.15 µM of HSP90-CDC37-BRAF^V600E^ complex was mixed with 0.3 µM of PP5 in a buffer containing 100mM NaCl, 25mM Hepes pH 8, 10mM KCl, 1mM MnCl2, 0.2mM TCEP, 2% glycerol and 2.5mM MgCl2. The reaction was started by incubating the samples at 30°C. Samples were taken over 45 minutes for SDS-PAGE analysis. The phosphorylation state of CDC37 Ser13 was probed by Western blot using a phospho-Ser13 specific antibody (Sigma).

### Mass Spectrometry Phosphorylation Analysis

Samples as in the dephosphoryation assays (above) of HSP90-CDC37-BRAF^V600E^ were either untreated or incubated with PP5 as above, and then split in two equal parts and diluted up to 100 μL with 100 mM triethylammonium bicarbonate (TEAB) followed by one-step reduction/alkylation with 5 mM TCEP and 10 mM iodoacetamide for 45 min at room temperature. Proteins were then digested overnight with 50 ng/μL trypsin (Pierce). Peptides were labelled with the TMT-10plex reagents (four labels used) according to manufacturer’s instructions (Thermo) followed by C18 clean-up using the Pierce Peptide Desalting Spin Columns. Phosphopeptides were enriched with the High-Select(tm) Fe-NTA Phosphopeptide Enrichment Kit (Thermo). Both the enrichment eluent and flowthrough (FT) were further subjected to mass spectrometry analysis.

LC-MS analysis was performed on the Dionex UltiMate 3000 UHPLC system coupled with the Orbitrap Lumos Mass Spectrometer (Thermo Scientific). Each sample was reconstituted in 30 μL 0.1% formic acid and 15 μL were loaded to the Acclaim PepMap 100, 100 μm × 2 cm C18, 5 μm trapping column at 10 μL/min flow rate of 0.1% formic acid loading buffer.

Peptides were analysed with an Acclaim PepMap (75 μm × 50 cm, 2 μm, 100 Å) C18 capillary column connected to a stainless-steel emitter with integrated liquid junction (cat# PSSELJ, MSWIL) fitted on a PSS2 adapter (MSWIL) on the EASY-Spray source at 45 °C. Mobile phase A was 0.1% formic acid and mobile phase B was 80% acetonitrile, 0.1% formic acid. The gradient separation method at flow rate 300 nL/min was the following: for 65 min (or 95 min for FT) gradient from 5%-38% B, for 5 min up to 95% B, for 5 min isocratic at 95% B, re-equilibration to 5% B in 5 min, for 10 min isocratic at 5% B. Each sample was injected twice. Precursors between 375-1,500 m/z were selected at 120,000 resolution in the top speed mode in 3 sec and were isolated for HCD fragmentation (collision energy 38%) with quadrupole isolation width 0.7 Th, Orbitrap detection at 50,000 resolution (or 30,000 for FT sample), max IT 100 ms (or 50 ms for FT) and AGC 1×10^5^. Targeted MS precursors were dynamically excluded for further isolation and activation for 30 or 45 sec seconds with 7 ppm mass tolerance.

The raw files were processed in Proteome Discoverer 2.4 (Thermo Scientific) with the SequestHT search engine for peptide identification and quantification. The precursor and fragment ion mass tolerances were 20 ppm and 0.02 Da respectively. Spectra were searched for fully tryptic peptides with maximum 2 miss-cleavages. TMT6plex at N-terminus/K and Carbamidomethyl at C were selected as static modifications. Oxidation of methionine, deamidation of asparagine/glutamine and phosphorylation of serine/threonine/tyrosine were selected as dynamic modifications. Spectra were searched against reviewed UniProt human protein entries, peptide confidence was estimated with the Percolator node and peptides were filtered at q-value<0.01 based on decoy database search. The reporter ion quantifier node included a TMT quantification method with an integration window tolerance of 15 ppm. Only peptides with average reporter signal-to-noise>3 were used, and phosphorylation localization probabilities were estimated with the IMP-ptmRS node.

Statistical analysis was performed in Perseus software. Data have been deposited in the Protein Identification Database PRIDE with accession code :.

## Notes

### Competing Interest Statement

The authors have declared no competing interest.

