## Supplementary Table1 and Figures for "HSP90-CDC37-PP5 forms a structural platform for kinase dephosphorylation"

### SUPPLEMENTARY TABLE 1 – CryoEM data collection and model refinement parameters

| | | | |
|---|---|---|---|
| HSP90-CDC37-BRAF$^{V600E}$ (**HCK**)<br>HSP90-CDC37-BRAF$^{V600E}$-PP5 open (**HCKPo**)<br>HSP90-CDC37-BRAF$^{V600E}$-PP5 closed (**HCKPc**) | | | |
| Magnification | 96,000 | | |
| Voltage (kV) | 300 | | |
| Electron exposure (e-/Å$^2$) | 45.0 | | |
| Defocus range (μm) | -1.3 to -2.5 | | |
| Pixel size (Å) | 0.86 | | |
| Symmetry imposed | C1 | | |
| Initial particle images (no.) | 1,126,606 (before split into three models) | | |
| | **HCK** | **HCKPo** | **HCKPc** |
| Final particle images (no.) | 400,624 | 67,985 | 105,063 |
| Map resolution (Å) | 3.4 | 4.2 | 3.9 |
| FSC threshold | 0.143 | 0.143 | 0.143 |
| **Refinement** | | | |
| Initial model used (PDB code) | 5FWK, 1UWH | 5FWK, 1UWH, 1WAO, 5HPE, 1A17 | 5FWK, 1UWH, 1WAO, 5HPE, 1A17 |
| Map resolution range (Å) | 3.29 – 6.9 | 3.68 – 10.39 | 4.02 – 11.75 |
| Map sharpening B factor (Å) | -130 | -84 | -180 |
| Non-hydrogen atoms | 14052 | 17742 | 17560 |
| Protein residues | 1716 | 2173 | 2150 |
| Co-factor residues | 2 | 2 | 2 |
| **B-factors (Å$^2$)** | | | |
| Protein | 61.6 | 115.2 | 90.0 |
| Co-factor | 48.6 | 60.2 | 41.8 |
| **R.m.s. deviations** | | | |
| Bond lengths (Å) | 0.007 | 0.003 | 0.009 |
| Bond angles (˚) | 0.995 | 0.674 | 1.017 |
| **Validation** | | | |
| MolProbity Score | 2.1 | 2.0 | 2.1 |
| Clashscore | 10.5 | 10.1 | 9.6 |
| Poor rotamers (%) | 0.0 | 0.1 | 0.3 |
| **Ramachandran plot** | | | |
| Favoured (%) | 89.3 | 92.0 | 90.3 |
| Allowed (%) | 10.1 | 7.8 | 9.1 |
| Disallowed (%) | 0.5 | 0.2 | 0.6 |

### SUPPLEMENTARY FIGURE 1 - PP5 activity and complex assembly

A

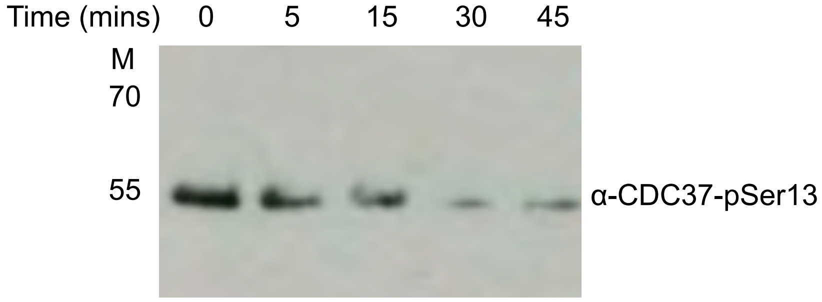


Western blot developed with phosphospecific antibody to pSer13 showing time course of dephosphorylation of CDC37-pSer13 when PP5 is added to an HSP90-CDC37-BRAF$^{V600E}$ complex

B

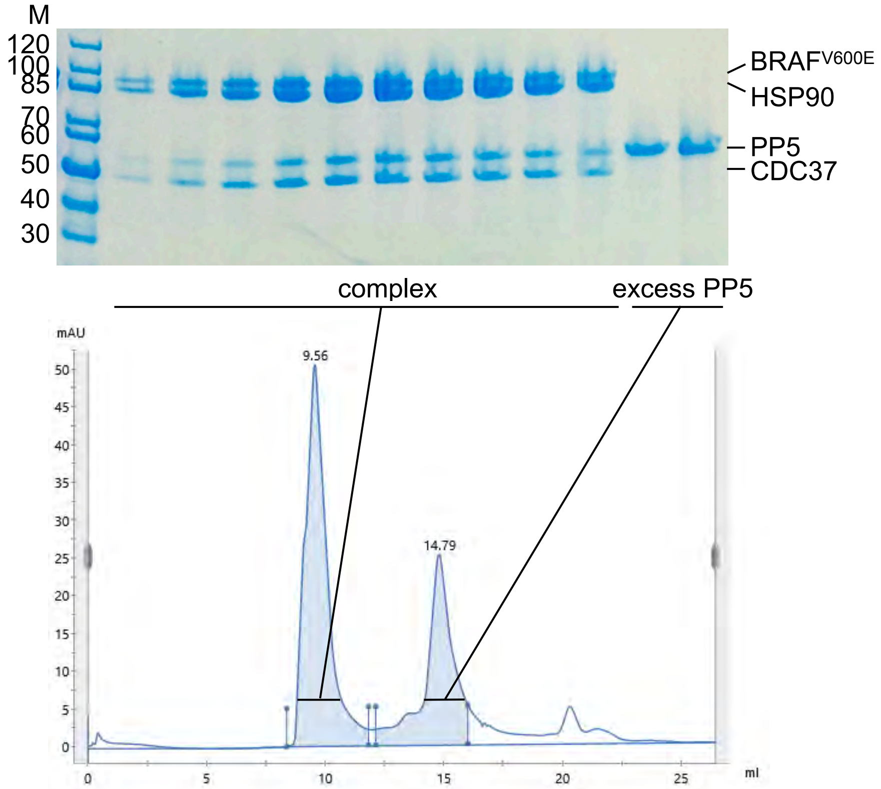


Coomassie-stained SDS PAGE gel showing purification of HSP90-CDC37-BRAF$^{V600E}$-PP5 complex by size exclusion chromatography. Elution profile from S200 is shown below.

C

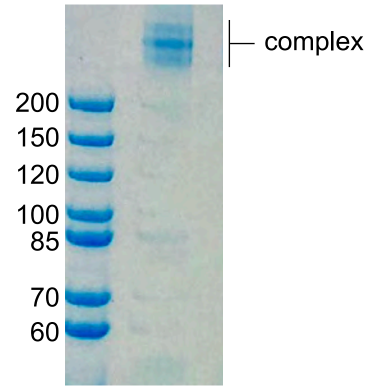


Coomassie-stained SDS PAGE gel showing high molecular weight complex formed by BS3 cross linking of peak fractions from B

D

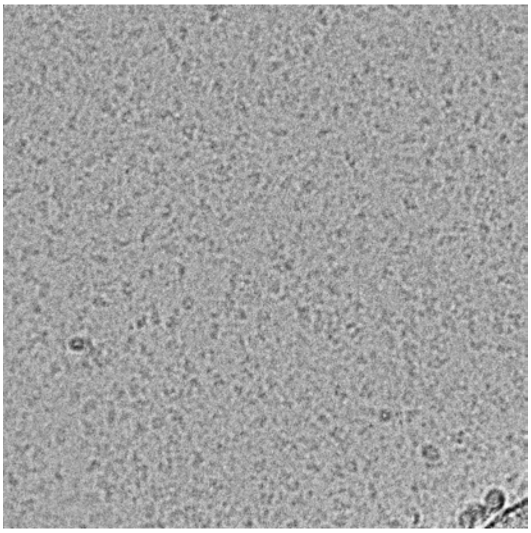

Example electron micrograph of cryogenically preserved sample of C applied to grids,

E

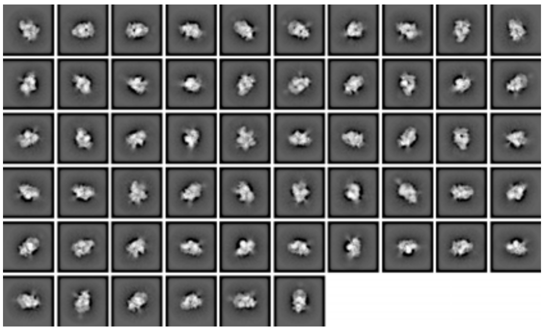

2D class averages of particles picked from micrographs, with inverted contrast. Clear secondary structure elements are visible at this stage.

### SUPPLEMENTARY FIGURE 2 - CryoEM processing schema

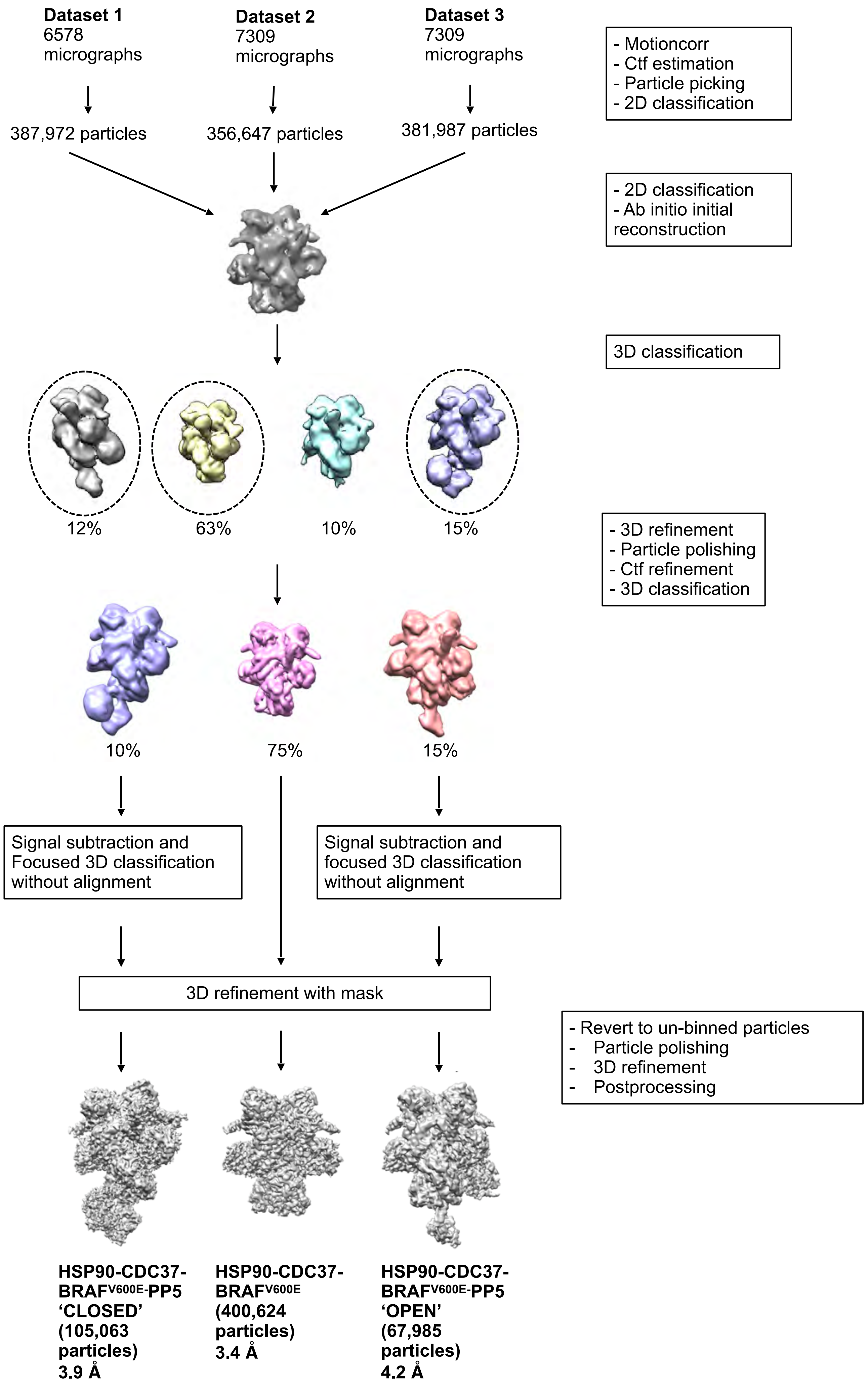

### SUPPLEMENTARY FIGURE 3 - cryoEM resolution

HSP90-CDC37-$BRAF^{V600E}$

HSP90-CDC37-$BRAF^{V600E}$ -PP5 closed

HSP90-CDC37-$BRAF^{V600E}$ -PP5 open

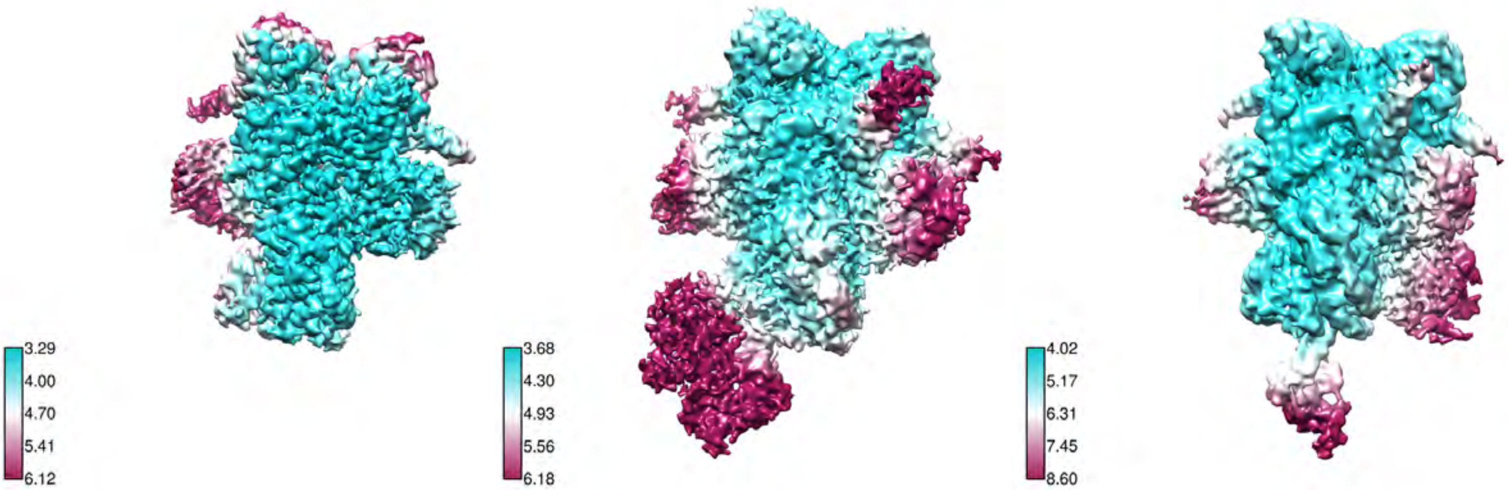


CryoEM volumes for the three structures reported here, surface coloured to reflect the local resolution estimated by RELION4.0

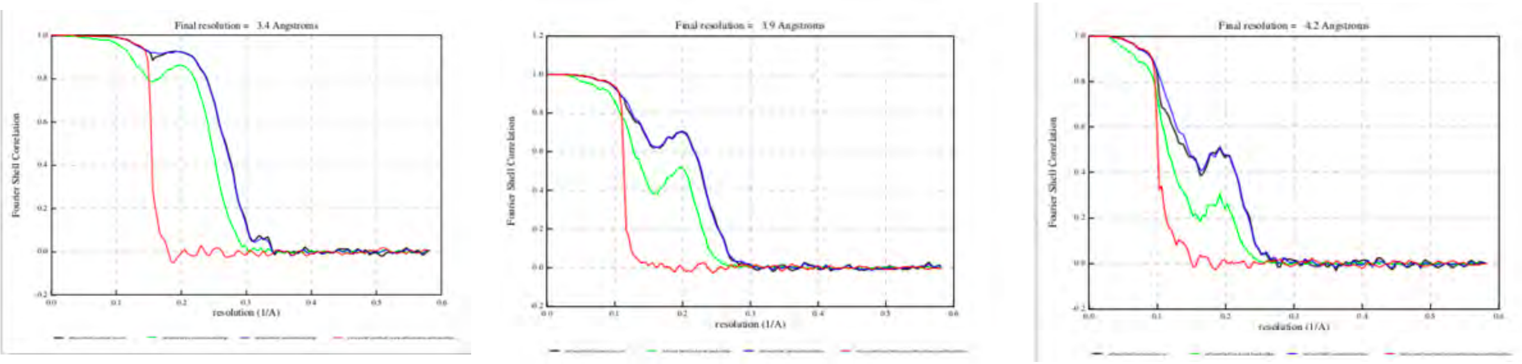


Fourier Shell Correlation plots for the three structure reported here.

### SUPPLEMENTARY FIGURE 4 - PP5 target sites in BRAF$^{V600E}$

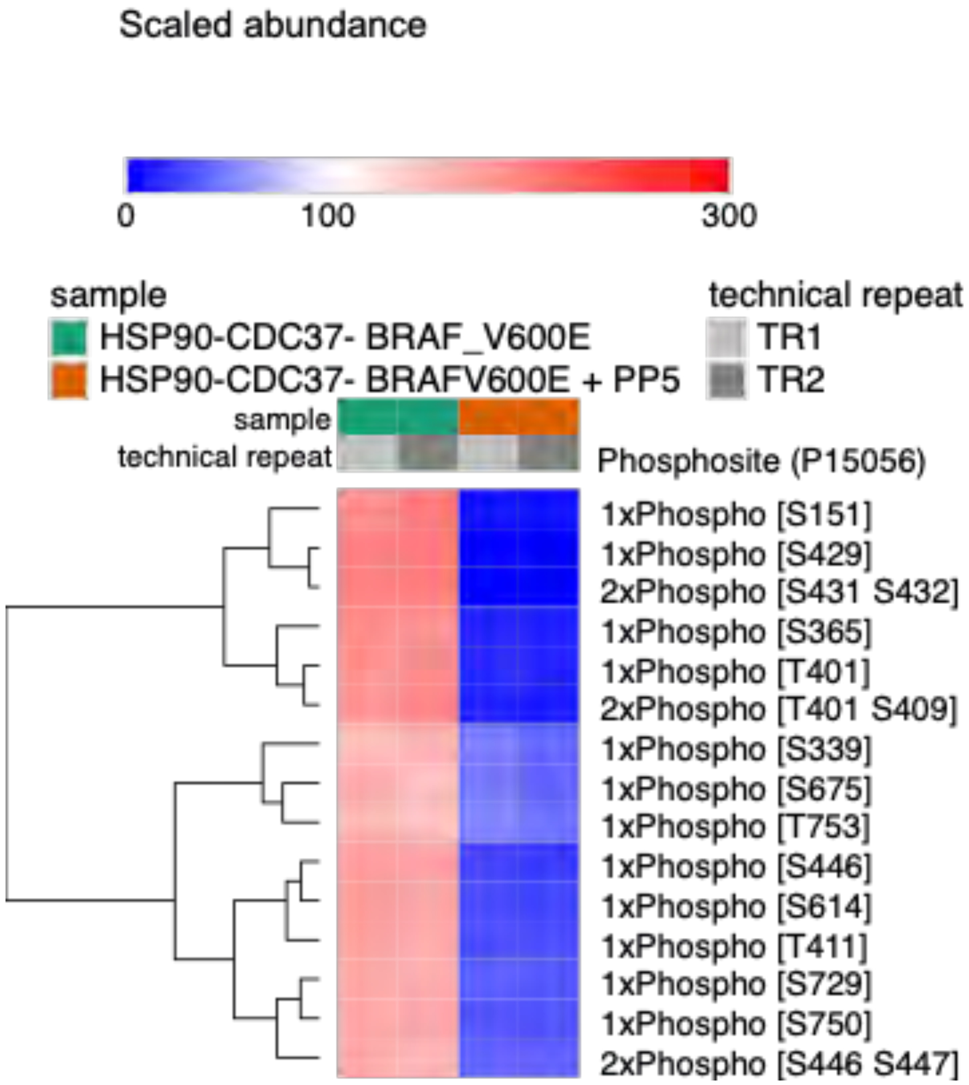


Comparison of phosphopeptide abundances from mass spectrometry analysis of BRAF$^{V600E}$ in the context of a complex with HSP90-CDC37, with or without incubation with PP5, each with two technical repeats utilising different labelling. Peptides are clustered on scaled abundance. The full dataset is provided as a Supplementary file.
